# The Rise of Polyploids During Environmental Upheaval

**DOI:** 10.1101/2024.11.22.624806

**Authors:** Hengchi Chen, Fabricio Almeida-Silva, Garben Logghe, Steven Maere, Dries Bonte, Yves Van de Peer

## Abstract

Polyploidy, or whole-genome duplication (WGD), serves as both a significant evolutionary force and a potential evolutionary dead end, occurring extensively across the tree of life, particularly among angiosperms. Despite the prevalence of polyploid organisms, instances of ancient polyploidy are surprisingly rare, presenting a paradox that remains poorly understood. In this study, we constructed a comprehensive genomic dataset of 470 angiosperm species to address this issue. We developed a highly congruent evolutionary timescale and dated 132 ancient WGD events that are non-randomly distributed, revealing a clustering around pivotal periods of environmental upheaval and extinction. Notably, our findings highlight a strong correlation between waves of paleopolyploidization and significant events such as the Middle Miocene Disruption, the Eocene–Oligocene transition (EOT), the Paleocene–Eocene Thermal Maximum (PETM), the Cretaceous–Paleogene (K–Pg) extinction, and different oceanic anoxic (OAE) events, several of which can be linked to extinction events impacting flowering plant genera. By integrating multiple lines of evidence, we propose that polyploid organisms have an increased chance of survival during times of great environmental turmoil, a conclusion with important implications in the context of contemporary climate change and rapid global warming.

## INTRODUCTION

The realization that the evolutionary trajectory of most, if not all, flowering plants has been profoundly influenced by multiple rounds of polyploidization - the result of whole genome duplication (WGD) - has been a cornerstone of early plant genomics research^1^. These "remarkable cycles of polyploidy" had been predicted by Haldane (1933)^2^ in the early days of polyploidy research, but were later often considered as “evolutionary noise”^3^ or even as evolutionary “dead ends”^4,5^. Such skepticism seemed partially justified: although extant polyploid organisms significantly enrich botanical diversity^6^, and polyploidization is acknowledged as a major driver of plant speciation^7^, the long-term persistence of polyploids appears to be a rare and infrequent phenomenon^8^. This infrequency suggests that substantial numbers of polyploid lineages may falter, failing to endure through evolutionary timescales. The presence of numerous instances of recurring polyploidy and recently originated polyploids, against a backdrop of scarce evidence for ancient polyploidy, presents an interesting evolutionary paradox^8,9^.

Whole genome duplication carries transformative implications as a significant mutational occurrence. WGD can lead to immediate detrimental fitness effects, including complications in cell cycling, stunted growth, and reproductive incompatibility with ancestral (diploid) forms. Simultaneously, WGDs elicit an instantaneous increase in cell dimensions, various genomic and epigenomic alterations, along with an array of downstream effects that directly influence fitness^10^. Both disadvantages and potential benefits of WGDs have been extensively documented^11-13^. Although the long-term consequences of WGD are relatively easy to understand (i.e., extra genetic material created by WGD can lead to novel gene functions and more robust regulatory networks^14^), the immediate or short-term advantages following WGD are less apparent. Several recent studies have meticulously described the immediate effects of polyploidy^12,15,16^. Among the most consistent outcomes of WGD is amplified cell size, coupled with physiological modifications that can lead to increased resilience in stress conditions. For instance, neo-autotetraploid cultivars of *Arabidopsis thaliana* display increased tolerance to salt and drought stress compared to their diploid progenitors^17,18^. Similarly, tetraploid cultivars of rice (*Oryza sativa*) and citrange (*Citrus sinensis* × *Poncirus trifoliata*) exhibit increased tolerance to salt and drought stress due to WGD’s modulation of genes involved in stress response and phytohormone signaling pathways^19,20^. As these examples of physiological and cellular responses to stress are frequently observed in cultivated polyploids, they have led to the hypothesis that polyploidization generally aligns with an increased resilience to stress^11,13^.

An enhanced stress tolerance or adaptive flexibility is hypothesized to transiently elevate polyploid establishment rates beyond expected norms during periods of environmental change. While classical birth–death models have successfully linked speciation rates to environmental variables like temperature^21,22^, validating similar hypotheses for polyploid establishment rates remains challenging because the environmental drivers and responses are highly diverse, likely often interacting. In 2009, Fawcett and co-authors^23^ estimated the absolute ages of nine ancient WGD events in angiosperms and found that these events appeared to roughly coincide with the K–Pg boundary, based on which they hypothesized that polyploidization may have helped plant lineages survive the K-Pg mass extinction event. Some of the same authors^24^ expanded the list of nine WGDs with another 11 and confirmed a clustering around the K–Pg boundary. These two studies, although invigorating a range of hypotheses about the evolutionary impact of polyploidy^13,25-27^, were not only constrained by limited phylogenetic sampling, but also remained contentious in light of recent large-scale WGD studies, where a clustering of WGDs in time, or a link with environmental disturbance, was not observed or reported^26-28^. Moreover, the inference and dating of ancient WGDs is notoriously difficult^29^ and numerous previously identified ancient WGD events have been shown to be either false positives^30,31^, or wrongly dated, or both^32,33^.

Here, we conducted an extensive analysis of a dataset encompassing 470 angiosperm genomes to identify and date 132 independent ancient polyploidization events using the Bayesian relaxed molecular clock model and 44 fossil calibrations for angiosperms that were highly curated following best practices^34^. To investigate patterns of paleopolyploidization across lineages in conjunction with paleoclimate dynamics and periods of mass extinction, we first meticulously assembled a robust phylogenetic framework and evolutionary timeline for the angiosperms. Our findings show that paleopolyploidization occurs across each major clade of angiosperms, including the ANA grade, Ceratophyllales, Chloranthales, eudicots, monocots, and magnoliids, and spans an evolutionary timeline from the Lower Cretaceous to the Neogene. Although the frequency of paleopolyploidization across different lineages and time varies dramatically, we find that ancient WGD establishment is correlated with periods of environmental turmoil or mass extinction, and more generally with periods of reduced lineage diversity and associated reduced competition. To facilitate data reuse, we built a web application (available at https://bioinformatics.psb.ugent.be/AngioWGD/, Supplementary Note 1) where users can explore WGDs across the angiosperm phylogeny, download inferred dates for each WGD, and access original genomic data included in this work.

## RESULTS

### Genome assessment based on BUSCO and OMArk

Because the quality of genome assemblies is essential for reliable downstream comparative analyses, we assessed all genome assemblies, i.e., 470 angiosperm genomes and the genome of the outgroup species *Cycas panzhihuaensis*. Complete BUSCO^35^ scores are generally high (median, Q1, and Q3 = 92%, 85.15%, and 96.45%, respectively), and so are OMArk^36^ scores (median, Q1, and Q3 = 95.45%, 90.41%, and 96.52%, respectively) (Figures S1 and S2 in Supplementary Note 2). The percentages of missing BUSCOs are low (mean and median = 7.93% and 4.10%, respectively), and so are percentages of OMArk missing families (mean and median = 7.85% and 5.55%, respectively). Taken together, these statistics indicate the high quality of our datasets.

### Reconstruction of an evolutionary timescale of angiosperms

Using 32 gene tree datasets, each with 1,614 reference single-copy gene families obtained from the *embryophyta_odb10* dataset^37^, we inferred species trees and assessed their overall accuracy and consistency, and ultimately reconstructed a ‘consensus tree’ (see Method details). The resulting consensus tree (hereafter referred to as ‘main tree’), covering 45 orders, 131 families, and all major clades of Mesangiospermae and ANA grade, displays high dataset- and branch-wise concordance (Figure S2 and S3-7 in Supplementary Note 3). High congruence was achieved across alternative Multiple Sequence Alignment (MSA) datasets, while two discordances pertaining to Chloranthales and Crossosomatales were further tested, concluding that the phylogenetic relationships that Chloranthales is sister to magnoliids, and Crossosomatales is sister to Fabales and Myrtales, were most supported (Figure S3 and S5 in Supplementary Note 3). We also show that our phylogenetic inference is robust to random sampling bias of gene trees (Figure S7 in Supplementary Note 3).

In comparison to a recent large-scale angiosperm phylogeny based on 353 nuclear orthologues^38^, our main tree shows concordance in most of the ordinal-level relationships (Figure S8 in Supplementary Note 3), and in particular the results of our statistical topology tests on Chloranthales and Crossosomatales are supported. However, the phylogenetic positions of magnoliids, Ceratophyllales, and Caryophyllales remain uncertain. Although the position of magnoliids has long been disputed, several recent large-scale phylogenetic studies^26,39,40^ agree with our main tree, supporting monocots being sister to magnoliids and eudicots. Phylogenetic inference from 30 out of 32 datasets supports our main tree, while the only two datasets that did not (the codon-level and TAPER-masked^41^ datasets at the branch filtering cut-off of 70%) also fail to recover monophyly of several orders and denoted numerous branches as highly uncertain (Supplementary Note 3), probably due to an insufficient number of remaining branches after stringent filtering. The phylogenetic location of Ceratophyllales in Zuntini et al. (2024)^38^ is highly uncertain (local posterior probability (LPP) below 0.5), and it was claimed by the authors themselves as inconclusive, while our main tree fully supports Ceratophyllales as sister to the eudicots (LPP = 1). Our tree also supports Caryophyllales as sister to the Lamiids and Asterids (LPP = 0.83) in 31 out of 32 datasets (except for the codon-level dataset at the branch filtering cut-off of 70%), in line with previous studies^26,39,40,42^, but disagreeing with Zuntini and co-authors^38^, who showed high gene tree conflict at the node pertaining to Caryophyllales.

Using the topology of our main tree, we reconstructed a time-calibrated phylogenetic tree based on our angiosperm fossil calibration dataset (Supplementary Note 4). Given that only 1 Single-Copy Orthogroup (SOG, see Method details) exist including all 470 angiosperm species, we slightly relaxed the species coverage to 99%, resulting in 12 SOGs encompassing 466 species. The uncorrelated lognormal (UCLN) relaxed molecular clock model was applied with soft fossil calibration bounds to consider both rate uncertainty and time uncertainty (Supplementary Note 4), resulting in an evolutionary time tree of 466 angiosperm species (Figure 1). The stem order and family ages in our time tree (Table S3.2 and 3.3) are highly correlated with those obtained by Ramírez-Barahona and co-authors^43^ (Pearson correlation coefficients = 0.8157 and 0.7542; P <0.0001; Figure S9 in Supplementary Note 5), suggesting comparable divergence time estimates.

**Figure 1.**
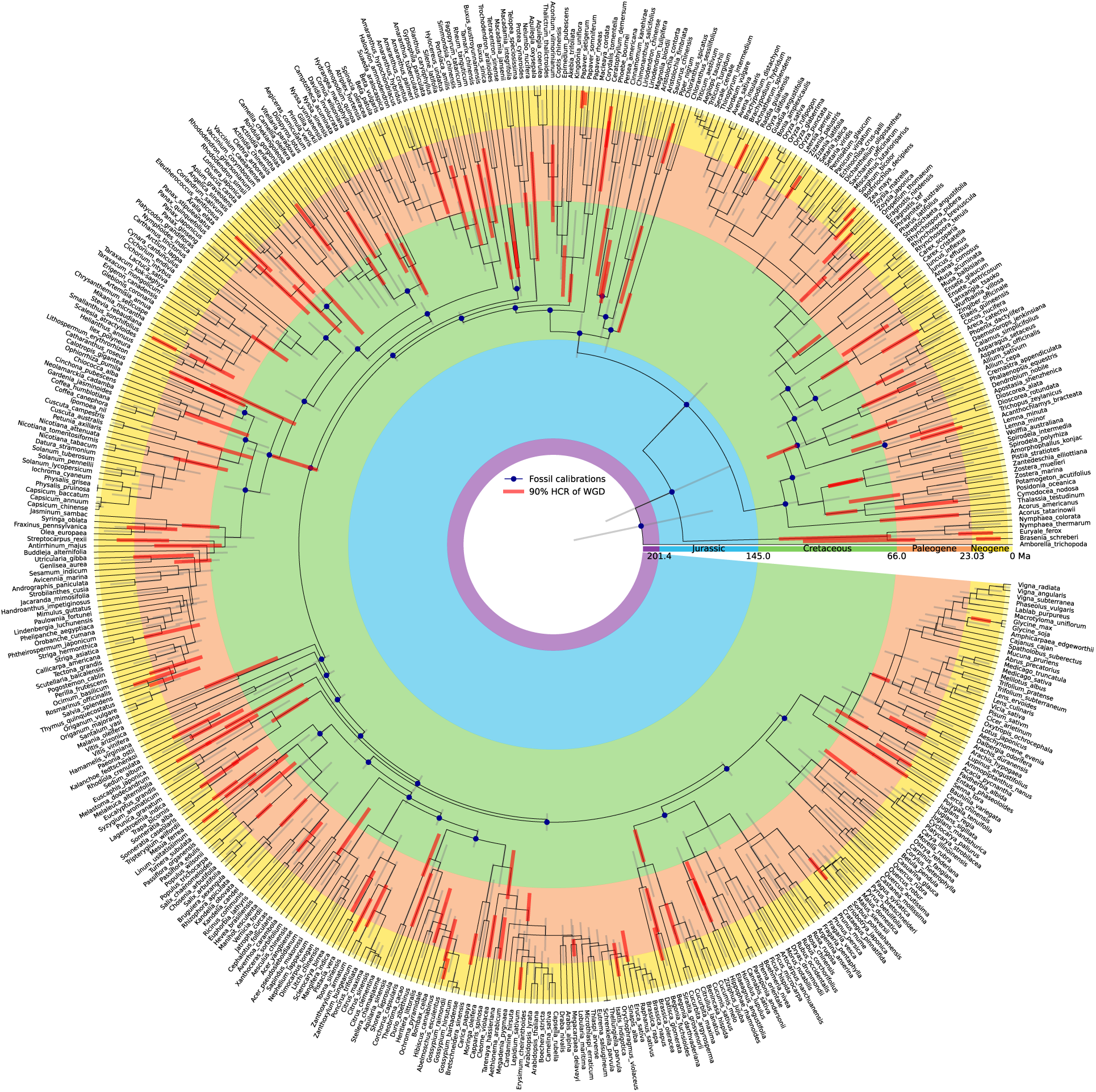
Reconstructed evolutionary time tree of 466 angiosperms. The 95% Highest Posterior Density (HPD) of divergence time estimates are represented as light gray lines on the internodes, while the 90% Highest Convergence Region (HCR) (see Methods) of WGD date estimates are represented as red rectangles. Applied fossil calibrations are represented as dark blue dots. Geological epochs are marked in distinct colors.

### Dating WGDs reveals temporal and phylogenetic dynamics of paleopolyploidization

We identified 132 independent ancient WGD events including both previously reported and novel ones, and conducted standardized WGD dating analysis on each one of them (Figure 1; Supplementary Note 6). Eudicots display the largest number of ancient WGD events (*N* = 95), followed by monocots (*N = 25*), magnoliids (*N* = 5), ANA grade (*N* = 4), Ceratophyllales (*N* = 2), and Chloranthales (*N* = 1). WGD frequency among different orders and families varies greatly. For instance, Lamiales harbors 10 independent ancient WGD events, which is greater than the 95^th^ percentile of all orders with ancient WGDs (Figure S1b in Document S1; Figure 2). Likewise, the plant families Poaceae, Fabaceae, and Brassicaceae contain 8, 5, and 5 independent ancient WGD events, respectively, which is greater than the 95^th^ percentile of all families with ancient WGDs (Figure S1c in Document S1; Figure 2).

**Figure 2.**
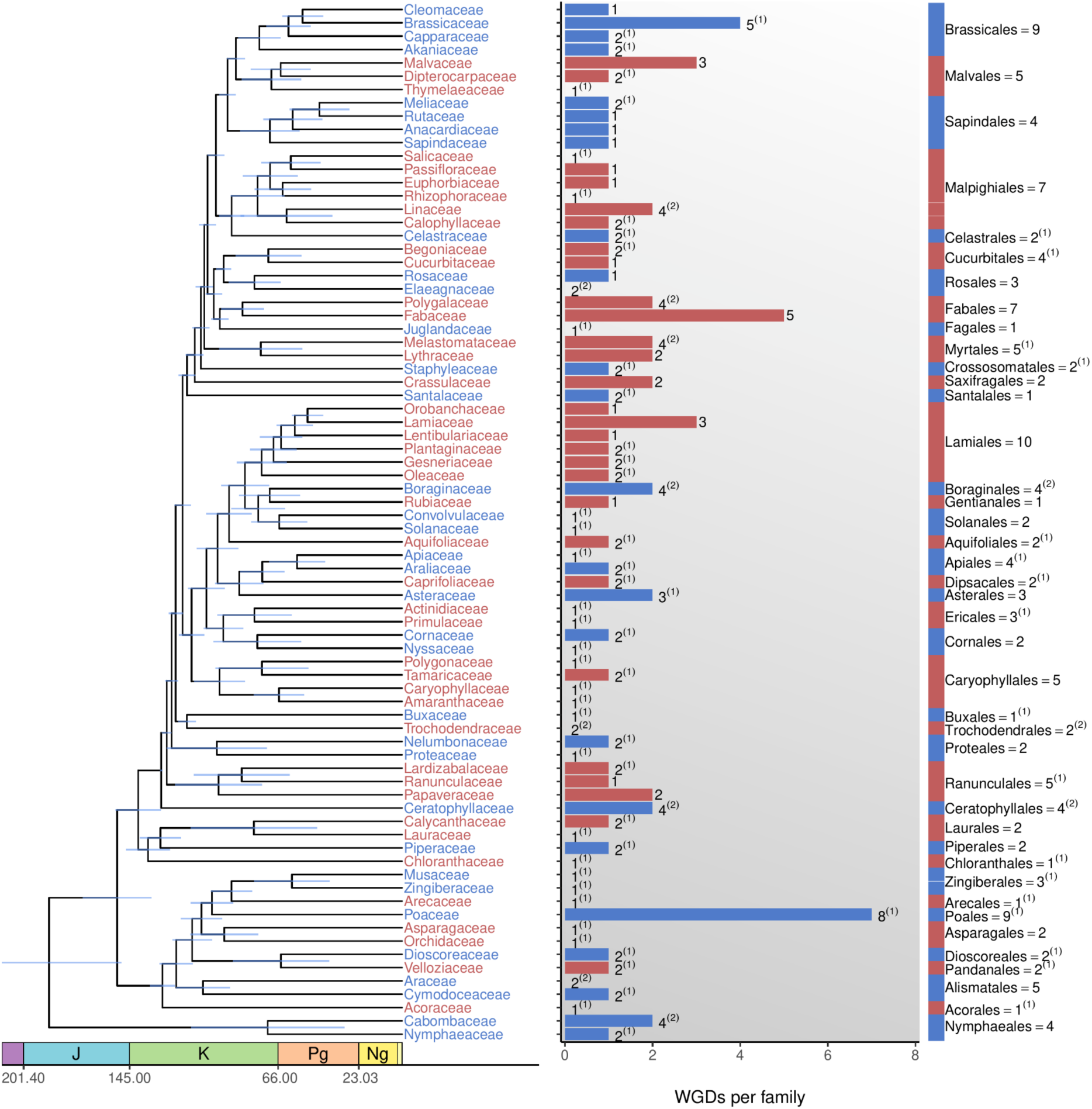
Distribution and number of WGDs in the plant families and orders considered in the current study. Numbers indicate WGDs observed in the clade. Numbers in superscript in parentheses indicate the number of WGDs at the base of each clade (MRCA), i.e., shared by all descendants. For instance, ‘Myrtales = 5^(1)^ indicates five WGDs in Myrtales, one of which occurred in the most recent common ancestor of this order.

The most ancient WGD event dated in monocots is the well-known τ event^44^, shared by all non-Alismatales monocots (excluding Acorales) and dated at 121.31 Ma (90% HCR = 112.87-129.35 Ma) (Figure S1d in Document S1; Supplementary Note 6). In eudicots, the oldest dated WGD events are the ‘RANU’ event (WGD nomenclature in Supplementary Note 6), shared by Ranunculales (Figure S1e in Document S1) and dated at 113.92 Ma (90% HCR = 108.24-122.32 Ma), and the γ event^45^, shared by core eudicots (Figure S1d in Document S1) and dated at 107.58 Ma (90% HCR = 94.96-119.90 Ma). In magnoliids, the oldest WGD event is the ‘LAUR β’ event, shared by Laurales and Magnoliales (Figure S1d in Document S1) and dated at 109.98 Ma (90% HCR = 98.56-120.51 Ma). There are 10 independent ancient WGD events shared by entire plant orders, and 10 ancient WGD events shared by ‘partial’ orders (i.e., shared by multiple plant families, but not the entire order; Figure S1e and S1f in Document S1). We also identified 31 independent ancient WGD events shared by entire plant families (Figure S1g in Document S1), and 20 ancient WGD events shared by ‘partial’ families (i.e., shared by multiple plant genera, but not the entire family; Figure S1h in Document S1). The remaining 58 independent paleopolyploidy events are shared by entire genera or unique to specific species (Figure S1i in Document S1).

### The WGD establishment rate is negatively correlated with species richness across time

Linear regression analysis revealed a strong linearity between the ancient WGD establishment rate (i.e., the number of ancient WGDs observed per lineage and per time unit) and the number of lineages across time (*R*^2^ varying from 0.2734 to 0.8217 when using different time window sizes, P<0.0001; Figure S3 in Document S1), and a linear model fit better than a power-law model (Figure S4 in Document S1). As the number of lineages increases exponentially in time (*R*^2^=0.9915, P<0.0001; Figure S5 in Document S1), the negative linear dependence of the WGD establishment rate on species richness translates to an exponential relationship between the WGD establishment rate and time.

### Significant deviations from the exponential relationship between WGD establishment rate and time unveil WGD clustering in time

When modeling the WGD establishment rate as a linear function of species richness and time and fitting a smoothing spline to the resulting relative residuals, 10 WGD peaks are recovered in the past 115 Ma (Figure 3A). An alternative approach using exponential-Gaussian mixture modeling of the WGD establishment rate profile with a varying number of Gaussians recovers all these peaks except the most recent one (around 8 Ma) in the optimal mixture model (adjusted *R*^2^=0.9987, Figure 3D, see Table S4). In addition, comparison of the observed peaks with a null distribution of the WGD establishment rate across time, as obtained from a stochastic WGD model assuming constant establishment rates in major clades, shows that 6 of the peaks rise above the 95% confidence interval (CI) of the null distribution (Figure 3G), namely those located at 35, 54, 64, 73, 85 and 99 Ma. These results suggest that at least 6 of the peaks recovered in the relative residuals profile may be evolutionarily significant. A null model including negative lineage diversity dependence of the WGD establishment rate showed two peaks at 73 Ma and 54 Ma beyond the 95% CI (Figure S6 in Document S1). The better model fit confirms again that a lineage diversity-dependent WGD establishment rate model is a better fit to the data than a constant rate model, supporting the hypothesis that the background WGD establishment rate is lineage diversity-dependent. For the two peaks that reach the upper 95% CI threshold, the lineage diversity-dependent null model provides solid evidence that other factors than random lineage diversity-dependent WGD establishment contributed to their formation. For the four peaks at 35, 64, 85, and 99 Ma that exceed the upper 50% CI threshold, this cannot be assessed with much confidence from this model, but one must consider that the null model WGD establishment rate parameters were fitted to all the data, including the peaks. If the observed peaks are non-random and superimposed on a random lineage diversity-dependent background, the background WGD establishment rates in the null model and the associated upper CI thresholds would have been overestimated.

**Figure 3.**
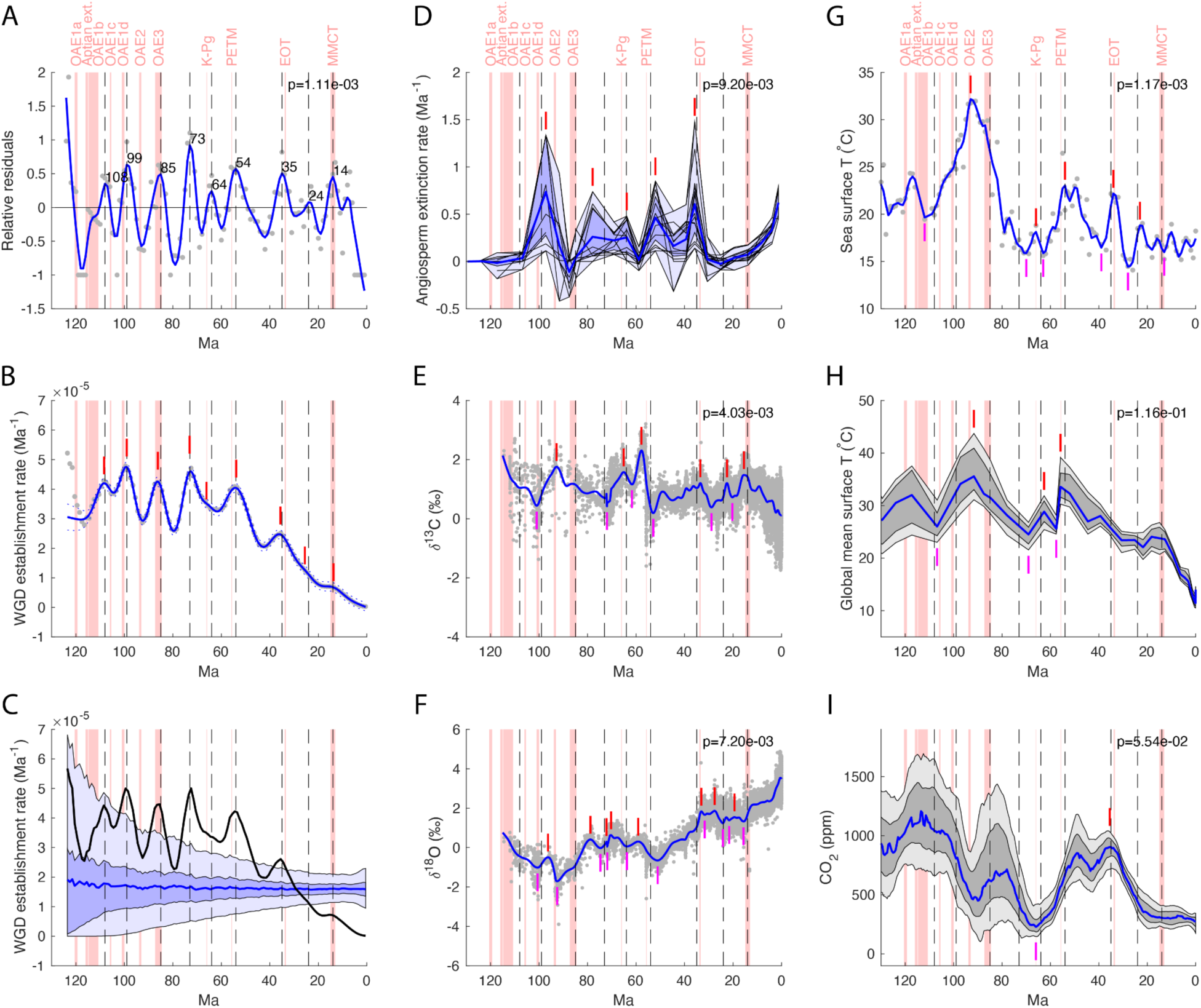
Association between WGD establishment rate peaks, geological events, extinction events and paleoclimatic extrema. Pink bars in all panels indicate significant extinction events or oceanic anoxic events (OAEs) in the past 130 Ma, as labeled in the top panels. Aptian ext.: Aptian extinction; K–Pg: Cretaceous–Paleogene extinction; PETM: Paleocene–Eocene Thermal Maximum; EOT: Eocene–Oligocene Transition; MMCT: Middle Miocene Climatic Transition. Vertical black dashed lines in all panels indicate WGD establishment rate peak locations in panel A. **A**. Gray dots represent the relative residuals of a linear model in which the moving average of the WGD establishment rate (window size 4 Ma) is modeled as a function of time and the number of angiosperm lineages in the 466-species tree through time (Figure 1). The blue curve is a LOESS fit with span set to 10% of the total number of data points. Peaks in the blue curve that are recovered in the optimal exponential-Gaussian mixture model (panel B) are indicated with vertical black dashed lines and the associated numbers indicate their ages in Ma. The extinction events and OAEs displayed in pink are located significantly closer to the WGD peaks than expected under a null model of uniform random WGD peak placement across the past 115 Ma, as indicated by the P-value in the upper right corner of the panel. **B**. Optimal exponential-Gaussian mixture model for WGD establishment rate data. Gray dots are kernel density estimates (KDE) of the WGD establishment rate (KDE bandwidth = 3.1). The blue line is the optimal exponential-Gaussian mixture fit (adjusted *R*^2^ = 0.9986) to the KDE data for the past 115 Ma, containing an exponential background and 9 Gaussian components of which the means are indicated by vertical red bars. **C**. Comparison of the observed KDE profile for WGD establishment rate (black line) with a null model assuming constant clade-specific WGD establishment rates (see Methods). The blue line represents the average WGD establishment rate profile obtained across 10,000 null model simulations, while the dark and light blue bands depict the 50% and 95% confidence intervals, respectively. Six of the nine peaks identified in panels A and B surpass the 95% confidence threshold. **D**. Genus-level angiosperm extinction rate profiles for the past 130 Ma. Black lines are extinction rate profiles obtained from the angiosperm fossil record in the Paleobiology database^113^ using three methods of sampling standardization and four macroevolutionary rate models in divDyn (v0.8.3)^114^. The blue line is the average estimated angiosperm extinction rate across the 12 models used, with dark blue and light blue bands depicting the interquartile range and total range of the values. Five extinction rate peaks are indicated with vertical red bars. These angiosperm extinction rate peaks are located significantly closer to a WGD peak in panel A (dashed vertical black lines) than expected under a null model of uniform random WGD peak placement across the past 115 Ma, as indicated by the P-value in the upper right corner of the panel. **E**. Benthic foraminiferal δ^13^C isotopic signature over the past 115 Ma, data from Friedrich et al. (2012)^115^. Gray dots are δ^13^C measurements, the blue line is a smoothing spline with smoothing parameter 0.05. **F**. Benthic foraminiferal δ^18^O isotopic signature over the past 115 Ma, data from Friedrich et al. (2012)^115^. Gray dots are δ^18^O measurements, the blue line is a smoothing spline with smoothing parameter 0.05. **G**. Low-latitude sea surface temperature in the past 130 Ma, data from Song et al. (2019)^116^. Gray dots represent mean sea surface temperature estimates, and the blue curve is a LOESS fit with the span set to 2% of the total number of data points. **H**. Global mean surface temperature in the past 130 Ma, data from Judd et al. (2024)^117^. The blue line represents the median estimate while the dark and light gray bands represent the 68% and 90% confidence intervals, respectively. **I**. Atmospheric CO_2_ concentration in the past 130 Ma, data from Foster et al. (2017)^118^. The dark blue solid curve represents the probability maximum while the dark and light gray bands represent the 68% and 95% confidence intervals, respectively. Vertical red and purple bars in panels E-I represent the maxima and minima of the blue lines, respectively, for which the linkage to the observed WGD peaks in panel A was the strongest (minimum P-value in top-right corner of panels). For each of the panels E-I, sets consisting of the top-1 up to the top-18 extrema were tested by comparing the observed average nearest distance to a WGD peak for any given set of extrema to a null distribution of average nearest distances obtained by uniformly sampling 9 WGD peaks across the past 115 Ma a million times. The same randomization procedure was used for panels A and D but there only the full set of geological events (panel A) or angiosperm extinction rate peaks (panel D) was tested.

### Significant coupling between ancient WGD establishment rate peaks and environmental upheaval

To assess the evolutionary significance of the nine peaks in WGD establishment rate (further referred to as WGD peaks) recovered in the linear and exponential-Gaussian mixture models, we compared the timing of these peaks to the timing of geological events, mass extinction events, and significant maxima or minima in time profiles of paleoclimatic variables (Figure 3). Monte Carlo randomization analyses of WGD peak timing uncovered that the observed WGD peaks in the past 115 Ma are located significantly closer to known extinction events and oceanic anoxic events (OAEs) than expected by chance (P=1.11e-03, Figure 3A). The observed peaks are also significantly linked to genus-level angiosperm extinction rate peaks inferred from the angiosperm fossil record, as documented in the Paleobiology database (P=9.20e-03, Figure 3D). We also assessed links between the observed WGD peaks and minima/maxima in smoothed time profiles of benthic foraminiferal δ^13^C and δ^18^O isotopic signatures over the past 115 Ma (Figure 3E, F), and time profiles of inferred sea surface temperature (Figure 3G), global mean surface temperature (Figure 3H) and atmospheric CO_2_ concentration (Figure 3I) over the past 115 Ma. Given that the paleoclimatic profiles contain large numbers of minima and maxima, we focused on the top-18 minima/maxima for each profile in terms of peak importance value (i.e., twice the number of WGD peaks) and compared randomized WGD peaks to the top-1, top-2 … top-18 minima/maxima for each (see Table S5.1). The minimum P-values obtained for each of the data layers and the corresponding top minima/maxima are displayed in Figure 3 and indicate significant coupling of the observed WGD peaks to important minima/maxima for all paleoclimatic variables at P<0.01, except global mean surface temperature (P=0.116) and CO_2_ concentration (P=0.0554). Because uniform randomization of peaks across the last 115 Ma may not produce sufficiently realistic inter-peak distance profiles (e.g., 9 random peaks may be located at 1,2,…,9 Ma whereas this is highly unlikely in smoothed WGD establishment rate profiles), we performed alternative randomization analyses where inter-peak distances were sampled from a Gaussian distribution (μ=11.57, σ=3.37 Ma) fit to the observed distances between the consecutive WGD peaks in Figure 3A. These randomization analyses are highly restrictive, as they essentially assume that peaks in establishment rate profiles are always located ∼10 Ma apart on average, whereas the observed regular WGD peak spacing they mimic may be caused by geological or biological rather than technical effects. It has been conjectured for instance that mass extinctions, associated geological events such as continental flood basalt eruptions and OAEs, and paleoclimatic events such as glaciations may happen at regular intervals, linked to astronomical cycles such as the Milankovitch Earth orbital eccentricity cycle and Galactic cycles^46^. Nevertheless, the restrictive randomization analyses still result in significant links (P<0.05) between the observed WGD peaks and geological events (P=6.62e-03), and minima/maxima in sea surface temperature (P=1.20e-02) and δ^13^C isotopic signatures (P=3.49e-02), whereas links between observed WGD peaks and angiosperm extinction rate peaks or minima/maxima in δ^18^O and CO_2_ are still significant at P<0.1 (Figure S7 in Document S1, full results in Table S5.2).

## DISCUSSION

### Waves of paleopolyploidizations during angiosperm evolution

Analyzing an extensive dataset containing 470 complete angiosperm genomes, we found evidence for 132 ancient polyploidization events. This seems a lot but if we consider individual evolutionary lineages, most plant lineages seem to contain remnants of only one, sometimes two, and in rare cases, three, four, or five WGD events (Figure 2). The contrast between numerous instances of recurring polyploidy and recently originated polyploids, and the few cases - within the same evolutionary lineage - of established ancient polyploidies over the past 150 million years (Figures 1 and 2), is striking^8,9^. This observation can only be understood when the high rate of polyploid emergence is countered by a reduced establishment rate at longer evolutionary time scales. Furthermore, one can wonder whether the sparsely distributed WGDs that survived in the long run got established at random times, or whether these WGDs survived because they occurred at decisive moments in evolution. As we show here, different model-based analyses support ‘waves’ of WGD establishment around 14, 24, 35, 54, 64, 73, 85, 99, and 108 Ma (Figure 3), many of which are associated with periods of mass extinction and major climate change.

### Ancient polyploidy correlates with periods of environmental upheaval

The first three WGD establishment waves (from more ancient to more recent) at ∼108, 99, and 85 Ma approximately coincide with the oceanic anoxic events OAE1c (∼106 Ma), OAE1d (∼101 Ma) and OAE3 (∼87 Ma), respectively^47^. Oceanic anoxic events are significant periods in Earth’s history when large portions of the ocean became depleted of oxygen, leading to widespread marine dysoxia or anoxia^48^. These events are often associated with climate shifts (thermal maxima), geological events such as increased large igneous province (LIP) volcanism and extinction events. OAE3 was the last and one of the longest of such events in the Cretaceous, associated with high temperatures and atmospheric CO_2_ levels (Figure 3G-I), but not with an extinction peak for angiosperm genera (Figure 3D). The comparatively small WGD wave at ∼108 Ma and the linked OAE1c event are associated with lower temperatures and high atmospheric CO_2_ levels (Figure 3G-I), but again not with a peak in angiosperm extinctions (Figure 3D). The WGD wave at ∼99 Ma and the close by OAE1d event on the other hand are associated with a major angiosperm extinction rate peak and substantial excursions in δ^13^C and δ^18^O isotopic signatures (Figure 3D-F). Interestingly, no WGD wave seems to be linked to OAE2, the most significant oceanic anoxic event in the Cretaceous associated with a mass extinction event at the Cenomanian–Turonian boundary (∼93.9 Ma)^49^. It is tempting to speculate that the observed WGD peak at ∼99 Ma and the angiosperm extinction rate peak at ∼97.2 Ma (Figure 3D) may be linked to OAE2 rather than OAE1d. Alternatively, the WGD peak around OAE1d may have reduced the need for additional angiosperm polyploidizations at the Cenomanian-Turonian boundary.

The fourth, and most prominent, wave of ancient WGDs occurred at ∼73 Ma. Interestingly, this wave is not located close to a major established extinction event or an OAE (Figure 3A), but it is located close to a peak in the genus-level angiosperm extinction rate at ∼78 Ma (Figure 3D, note that the temporal resolution of the angiosperm extinction rate profile and the associated extinction peaks is low). At the end of the Cretaceous (∼80–66 Ma), a period with cooling temperatures^50^ (Figure 3G,H) and decreasing CO_2_ levels (Figure 3I) after the preceding Cretaceous ‘hothouse’ period, overall plant diversity decreased substantially and the rise of the angiosperms temporarily stalled^51^. In Antarctica, angiosperm species diversity exhibited a pronounced dip around the Campanian-Maastrichtian boundary at ∼72.1 Ma^51^, very close to the inferred WGD wave’s location at ∼73 Ma. The Campanian–Maastrichtian Boundary Event (CMBE) is characterized by substantial negative and positive excursions in the δ^13^C and δ^18^O isotopic signatures, respectively (Figure 3E,F). What caused these excursions is still debated, with two of the competing hypotheses being the buildup of temporary ice sheets in Antarctica associated with a drop in sea levels^52^, or a major change in intermediate to deep water circulation caused by the global cooling trend^53^. Despite not being associated with a major extinction event, the presence of a pronounced WGD peak around 73 Ma suggests that the period around the Campanian–Maastrichtian boundary may have been more important for angiosperm evolution than previously realized. Alternatively, the long-term evolutionary success of some of the polyploidizations that happened ∼73 Ma may have been catalyzed by the increased potential of the affected lineages to weather the subsequent climatological events in this turbulent geological time period, such as the Middle Maastrichtian Event (∼69.5 Ma), the Late Maastrichtian Warming Event (∼66.3 Ma) associated with increased volcanic activity of the Deccan Traps^54^, and ultimately the K–Pg extinction event (∼66 Ma). This hypothesis of increased adaptive potential of species due to earlier polyploidizations may also help explain why the ancient WGD peak and angiosperm genus extinction rate peak at the K–Pg boundary itself (see below) is smaller in size than might be expected for a major mass extinction event.

The fifth WGD establishment wave, at ∼64 Ma, occurred close to the K–Pg boundary. The K–Pg mass extinction, ∼66 Ma, is the most recent of the ‘Big Five’ mass extinctions, best known for the extinction of the non-avian dinosaurs^55,56^. Previously, Fawcett and co-authors^23^ and Vanneste and coauthors^24^ already suggested a potential clustering of WGDs around the K–Pg boundary. Here, using a genomic data set twelve times larger (N = 470, with 132 ancient WGDs), we still find support for this observation, although the 64 Ma WGD peak recovered in Figure 3 A,B is substantially smaller than the peaks before and after it (73 and 55 Ma). The angiosperm extinction rate exhibits a peak at 66 Ma (Figure 3D), but it is the smallest of the five peaks recovered from the divDyn analyses (see Methods), suggesting that the K–Pg extinction event had a comparatively minor effect on the survival of angiosperm genera. At least one phylogenetic study points in the same direction^57^. Given the large uncertainties in ancient WGD dating, Fawcett and co-authors^23^ and Vanneste and co-authors^24^ assumed that all WGDs that were dated between 55 and 75 Ma were likely associated with the K–Pg boundary. Now, using a dataset with many more species and ancient WGDs, and hence enhanced resolution, the evidence suggests that the previously observed clustering of ancient WGDs at 55–75 Ma is in fact associated with three separate geological events: the Paleocene–Eocene Thermal Maximum (PETM, see below), the K–Pg boundary and the CMBE) of which K–Pg may have been the least consequential in terms of angiosperm genus extinction and ancient WGD establishment.

The sixth wave of WGDs at ∼54 Ma is located near the Paleocene–Eocene Thermal Maximum (PETM event), ∼55.8 Ma, a period characterized by a 5 to 8°C increase in global mean surface and sea surface temperatures (Figure 3G-H)^58-60^, and a concomitant significant negative excursion in δ^18^O in lake bed and ocean sediments (Figure 3F). The PETM event is considered one of the most extreme climate changes in Earth’s history, and was characterized by a massive release of δ^13^C-depleted carbon to the ocean–atmosphere system^61-63^, causing a prominent negative excursion in δ^13^C in e.g. benthic foraminifera from ocean sediment cores (Figure 3E). The causes of this massive carbon release are still heavily debated, but may include volcanic intrusions in carbon-rich sediments followed by massive thermogenic δ^13^C-depleted methane release^60,64^ or increased volcanism causing an initial phase of global warming, followed by subsequent warming-induced release of δ^13^C-depleted methane from submarine clathrates on continental margins^65^. The PETM had significant impacts on biodiversity and many species of both terrestrial and marine organisms became extinct, including angiosperms (Figure 3D), or experienced substantial shifts in their distribution. The PETM saw the flourishing of subtropical forests, with many lineages adapting to the warmer conditions^65,66^. Previously, Cai and coauthors^67^ also reported a clustering of ancient WGD events around the Paleocene–Eocene transition, based on the dating of 22 independent ancient WGD events from genomic and transcriptomic assemblies of 36 Malpighiales species.

The seventh WGD establishment wave at ∼35 Ma is located very close to the Eocene–Oligocene transition (EOT), associated with the Eocene–Oligocene extinction event or ‘Grande Coupure’, about 33.5 Ma, which was characterized by a rapid shift to a colder climate that led to the development of a permanent Antarctic ice sheet and significant environmental changes. The transition from the warm conditions of the Eocene to the cooler, more variable climates of the Oligocene is one of the most important climate shifts in Earth’s history. This shift is associated with a significant drop in global temperatures, because of both tectonic activities and changes in oceanic circulation patterns^68,69^. Indeed, the EOT and associated WGD wave correspond to a peak in δ^18^O indicative of a cooling event and increasing ice volume^70^ (Figure 3F). That the EOT and the associated WGD wave seem to correspond to a sea surface temperature peak (Figure 3G) rather than the transition to colder temperatures (between 32 and 30 Ma in Figure 3G) may be due to dating uncertainties. The EOT is also associated with a mass extinction event, mainly among marine species^71^, but also greatly affecting flowering plants (see Figure 3D). Indeed, on land, the cooler temperatures and changing habitats led to shifts in vegetation and the extinction of plants and animals adapted to the warmer, more humid conditions of the Eocene^72^. Overall, the EOT is considered a critical period in Earth’s history, marking significant changes in climate and ecology.

The eighth WGD establishment wave around 24 Ma does not coincide with a documented major extinction event or OAE (Figure 3A) or a peak in the angiosperm extinction rate (Figure 3B), but approximately coincides with the Oligocene-Miocene Transition (OMT), a transient global cooling event ∼ 23 Ma that is associated with significant Antarctic ice sheet expansion. However, the δ^13^C (Figure 3E) and sea surface temperature (Figure 3C) profiles exhibit a maximum around this time, and the δ^18^O a minimum (Figure 3H), which may reflect a warming event in the late Oligocene rather than the OMT^73^. The discrepancies between the estimated OMT age and the observed excursions in δ^13^C, δ^18^O and sea surface temperature profiles may be due to dating uncertainties. As the 24 Ma WGD peak is the least prominent of the WGD establishment rate peaks supported by mixture modeling (Figure 3B) and is not supported by our null model simulations (Figure 3C, Figure S6) and not strongly connected to a known mass extinction event or major climate transition, this is the WGD peak we are least confident about. Nevertheless, its timing fits nicely in a continuous series of 6 WGD peaks associated with successive geological boundaries, including, in succession, the WGD peak at ∼85 Ma close to the Coniacian-Santonian and Santonian-Campanian age boundaries in the Upper Cretaceous (∼86.3 and ∼83.6 Ma, respectively), the WGD peak at ∼73 Ma close to the Campanian-Maastrichtian age boundary at ∼72.1 Ma, the ∼64 Ma peak close to the Cretaceous-Paleogene (K-Pg) period boundary (∼66 Ma), the ∼54 Ma and ∼35 Ma WGD peaks close to the Paleocene-Eocene and Eocene-Oligocene epoch boundaries in the Paleogene (∼56 Ma and ∼33.9 Ma, respectively), and the ∼24 Ma peak close to the Paleogene-Neogene period boundary (which coincides with the OMT, ∼23 Ma). Of the three remaining WGD peaks, only one is close to a geological boundary, namely the WGD peak at ∼99 Ma close to the period boundary between the Lower and Upper Cretaceous (∼100.5 Ma).

The ninth and most recent wave of WGD establishment at ∼14 Ma coincides with the Middle Miocene Climatic Transition (MMCT), also known as the Middle Miocene disruption. The MMCT occurred approximately 13 to 15 Ma and marks a significant shift from a warmer climate to a cooler one, leading to substantial environmental changes greatly affecting ecosystems and biodiversity^74,75^. As the climate cooled, forests began to decline in many regions, giving way to grasslands and open savannas. This shift led to adaptations in flora and the evolution of new faunal communities, including grazers that thrived in the new grassland habitats^76^. The MMCT is sometimes considered a significant extinction event^77,78^, while others see it rather as a transformative period that significantly affected species distributions and set the groundwork for the ecosystems known in the later stages of the Miocene and into the Pleistocene^79,80^. The MMCT is not associated with a peak in the genus-level angiosperm extinction rate (Figure 3B), but does correspond with a drop in sea surface temperature (Figure 3C), a sharp drop in δ^13^C (Figure 3E), indicative of a decrease in vegetation abundance, and a sharp rise in δ^18^O (Figure 3H), indicative of a decrease in seawater temperature and/or growing ice sheets^81^.

### Revisiting the polyploid paradox

Nature is teeming with polyploid species that have extra sets of chromosomes. Yet, ancient WGDs that survive over the long term are surprisingly scarce (Figure 2). The apparent paucity of (established) ancient genome duplications and the existence of so many species that are currently polyploid provides a fascinating but puzzling paradox^8,82^. The rarity of polyploidy events that get ‘fixed’ on the long run seems to suggest that polyploidy is usually disadvantageous, and an evolutionary dead end. However, there is growing evidence that polyploidization can sometimes help overcome environmental change (stress)^13,83,84^. At decisive moments in evolution, as again shown here, polyploids seem to have had an adaptive advantage and likely outcompeted many of their diploid progenitors. Despite much research, the mechanistic underpinnings of why and how polyploids might be able to outcompete or outlive non-polyploids remain elusive.

Here, we demonstrate for the first time that WGD establishment is enhanced under conditions of low lineage diversity, both early on in angiosperm evolution and during periods of major environmental upheaval and mass extinction. The negative lineage diversity dependence we observe for the background WGD establishment rate is very similar to the well-known negative lineage diversity dependence of speciation rates. Indeed, empirical evidence suggests that diversification tends to decelerate as evolution progresses^21,85-87^. This slowdown is traditionally attributed to diversity-dependent mechanisms, where increasing species richness intensifies competition for resources and space, leading to niche saturation^88,89^. Recent studies highlight the role of environmental perturbations such as fluctuations in sea level, δ¹³C, CO₂ concentrations, and temperature, as additional drivers of diversification dynamics^22,90,91^. Very similar dynamics are observed here for WGD establishment rates, suggesting that, as for speciation, increased diversity reduces the likelihood of successful WGD establishment due to competitive exclusion. Conversely, in periods of reduced diversity and competition, e.g., when major environmental changes cause previously well-adapted species to go extinct and new niches to open up, WGD establishment rates may temporarily increase. In first instance, environmental stress may lead to an increase in polyploid formation due to increased unreduced gamete formation^92^. These polyploids may then be better able to adapt to the changed environment than their diploid progenitors, either due to immediate adaptive advantages obtained through polyploidization^93^ or heterosis, or through higher adaptability on the longer term^6,94^. Examples of the first are duplicates of RNA-binding proteins involved in cold responses that exhibit convergent retention across diverse angiosperm lineages following independent WGD events occurring around the K–Pg boundary^95^, or gene duplicate advantages in low temperature and darkness around the K–Pg boundary, when global cooling and darkness were the two main stresses^96^. To take advantage of potential higher long-term adaptability however, the polyploid lineage must be able to survive long enough to allow for any initial fitness disadvantages to be overcome^11,12^ and for adaptive advantages to arise, e.g. through genome fractionation and diploidization^97^ and duplicate gene divergence. Here, reduced diversity and reduced competition may be crucial in facilitating newly emerged polyploids to persist long enough to gain the upper hand^10,94,98^. In some cases, lag times of several million years have been identified between ancient WGD events and the subsequent diversification of lineages thought to have been enabled by beneficial traits obtained from those WGDs^99-101^. Such long lag times suggest that it may often take a lot of time for polyploids to evolve adaptive traits, which may be challenging in stable environments with high competition. Alternatively, an established (paleo)polyploid lineage may have had to wait for millions of years to diversify until the ecological opportunity presented itself in the form of severe environmental change. Both scenarios are not mutually exclusive. In the aftermath of environmental turmoil, as niches become filled due to radiation and competition increases again, the window of opportunity for polyploids to get established on the long term is expected to close again. In summary, our data indicates that WGDs are a dead-end under relatively stable conditions (hence their scarcity on the evolutionary tree) but could confer a selective advantage in times of severe environmental change, providing a potential explanation for the polyploid paradox.

## CONCLUSIONS

The advent of genomics made it clear that the significance of polyploidy extends across all eukaryotes, and that most, if not all, extant species (including our own) carry the signature of at least one ancient whole genome duplication (WGD) in their history^1,8,102-104^. Nevertheless, finding evidence for ancient WGD can be complicated^105^ and dating these events, going back tens to hundreds of millions of years ago, is notoriously difficult. Here, we assembled and assessed a large angiosperm genome dataset comprising 470 different angiosperm species and used state-of-the-art methods to identify and date 132 independent, ancient WGDs. Analyzing such a large number of species and WGDs is necessary to be able to draw reliable conclusions about potential correlations between WGDs and ‘decisive’ moments in evolution, which was a limitation of some previous studies based on much smaller numbers of species and WGDs^23,24^. Furthermore, our large dataset allows for a resolution that was impossible to achieve previously. Model-based analyses and simulations clearly reveal that the inferred WGD events are not randomly distributed but rather cluster around periods of environmental upheaval. We find significant clustering of WGDs around the Middle Miocene Disruption, the Eocene–Oligocene Extinction Event, the Paleocene–Eocene Thermal Maximum, the K–Pg boundary, and several oceanic anoxic events. Our study again provides support for, what we call “the polyploid paradox”: neopolyploids abound, while the long-term survival of polyploidy seems to be rare (Figure 2). We hypothesize that, in general, and under relatively stable environments, the disadvantages and the costs of being polyploid do not outweigh the sometimes-observed advantages such as increased stress resilience^12,13^. However, when the circumstances are such that the conditions to survive are extremely challenging, polyploids could be the ‘hopeful monsters’^93,106-109^ that can outcompete their diploid progenitors. In this respect, we feel that polyploidy can indeed often be considered an evolutionary dead end, at least when interpreted as ‘rarely successful’^110-112^, but that it can be an evolutionary and ecological force in (very) unstable times. In this respect it is interesting to speculate that, in the current Anthropocene, a time during which humans are having a substantial impact on our planet, several new ‘ancient’ polyploidy events might be in the making.

## RESOURCE AVAILABILITY

### Lead contact

Further information and requests for resources should be directed to and will be fulfilled by the lead contact, Yves Van de Peer.

### Materials availability

All results generated in this study are available from the lead contact upon request.

### Data and code availability

- The genomic data involved in this study have been deposited at figshare at https://doi.org/10.6084/m9.figshare.27011128.v1 and https://doi.org/10.6084/m9.figshare.27011134.v1.
- The posterior samples of WGD dates for each independent WGD event have been deposited at figshare at https://doi.org/10.6084/m9.figshare.27596856.v1 and the associated mixed *K*_S_ distribution figures have been deposited at figshare at https://doi.org/10.6084/m9.figshare.29553929.v1.
- The code for the genome assessment, phylogenetic inference, molecular dating, WGD identification and dating, Poisson process models of WGD simulation, leave-one-out cross-validation (loo-cv) of KDE bandwidth, residual analysis, mixture modeling analysis, coupling analysis, and the extinction rate estimation is available at https://github.com/heche-psb/Angiosperm_WGDs.
- Any additional information required to reanalyze the data reported in this paper is available from the lead contact upon request.

## ACKNOWLEDGEMENTS

H.C. acknowledges funding from the Research Foundation—Flanders (FWO) (No. 3G032219). Y.V.d.P. acknowledges funding from the European Research Council (ERC) under the European Union’s Horizon 2020 research and innovation program (No. 833522). Y.V.d.P. and F.A-S. acknowledge funding from Ghent University (Methusalem funding, BOF.MET.2021.0005.01).

## AUTHOR CONTRIBUTIONS

Y.V.d.P. and H.C. conceived and managed the project. H.C. collected the genomic data and conducted the assessment analysis. H.C. conducted the phylogenetic and molecular dating analysis. H.C. conducted the WGD inference and dating analysis. H.C. conducted the WGD simulation analyses. G.L., D.B. and F.A.S. conducted the Bayesian phylogenetic clustering analysis in the first version of the manuscript. F.A.S. constructed the website. S.M. conducted the mixture modeling analysis of the WGD establishment rate profile and the coupling analyses between WGD establishment rate peaks and other data layers. D.B. conducted the diversity dependence analysis of the ancient WGD establishment rate. H.C. conducted the loo-cv analysis and the extinction rate estimation analysis. H.C., F.A.S., D.B., S.M., and Y.V.d.P. wrote the manuscript. All authors read and approved the manuscript.

## SUPPLEMENTAL INFORMATION

**Document S1. Figures S1–S8**

**Table S1. Tukey’s HSD test of different datasets at different branch support filtering cut-offs, related to Figure 1**

**Table S2. Age constraints for the stem group of different angiosperm orders, related to Figure 1**

**Table S3.1. Functional descriptions of the 12 SOGs adopted in molecular dating, related to Figure 1**

**Table S3.2. Posterior mean, 95% HPD of stem age of angiosperm orders, related to Figure 1**

**Table S3.3. Posterior mean, 95% HPD of stem age of angiosperm families, related to Figure 1**

**Table S3.4–3.8. Tukey’s HSD test of the relative nucleotide and amino acid substitution rate, related to Figure 1**

**Table S4. Adjusted *R*^2^ for the Exponential-Gaussian mixture modeling, related to Figure 3**

**Table S5.1. P-values for the association test under the null hypothesis of random uniform peak distribution, related to Figure 3**

**Table S5.2. P-values for the association test under the null hypothesis of restricted distances between randomized WGD peaks, related to Figure 3**

**Table S6. *K*_S_ peak values, credible ranges and WGD dates for the absolute dating of each independent WGD, related to Figure 1**

**Table S7. WGD establishment rate parameter in the homogeneous Poisson process model, related to Figure 3**

**Supplementary Note 1. The *AngioWGD* web application, related to Figure 1**

**Supplementary Note 2. Assessment of genome assembly, related to Figure 1**

**Supplementary Note 3. Phylogeny reconstruction, related to Figure 1**

**Supplementary Note 4. Fossil calibrations, related to Figure 1**

**Supplementary Note 5. Evolutionary timeline and rate variation, related to Figure 1**

**Supplementary Note 6. WGD identification and dating, related to Figure 1**

**Supplementary Information 1. Results of species tree inference, related to Figure 1**

**Supplementary Information 2. Fossil calibration map, related to Figure 1**

## STAR★METHODS

### KEY RESOURCES TABLE

| REAGENT OR RESOURCE | SOURCE | IDENTIFIER |
| --- | --- | --- |
| Deposited data |  |  |
| Coding sequence of 470 angiosperm species | Public data, reassembled in this paper | https://doi.org/10.6084/m9.figshare.27011128.v1 |
| General Feature Format files of 470 angiosperm species | Public data, reassembled in this paper | https://doi.org/10.6084/m9.figshare.27011134.v1 |
| Posterior samples of WGD dates for 132 independent WGD events in angiosperms | This paper | https://doi.org/10.6084/m9.figshare.27596856.v1 |
| Software and algorithms |  |  |
| BUSCO v5.4.3 | Simão et al.<sup>35</sup> | https://busco.ezlab.org/ |
| OMArk | Nevers et al.<sup>36</sup> | https://omark.omabrowser.org/ |
| HMMER v3.1b2 | Potter et al.<sup>119</sup> | https://www.ebi.ac.uk/Tools/hmmer/ |
| MAFFT v7.310 | Katoh et al.<sup>120</sup> | https://mafft.cbrc.jp/ |
| TAPER | Zhang et al.<sup>41</sup> | https://github.com/chaoszhang/TAPER |
| IQTREE v1.6.5 | Nguyen et al.<sup>121</sup> | http://www.iqtree.org/ |
| ModelFinder | Kalyaanamoorthy et al.<sup>122</sup> | http://www.iqtree.org/ModelFinder/ |
| ASTRAL-Pro | Zhang et al.<sup>123</sup> | https://github.com/chaoszhang/ASTER |
| PAML v4.9j | Ziheng Yang<sup>124</sup> | http://abacus.gene.ucl.ac.uk/software/paml.html |
| i-ADHoRe v3.0.01 | Proost et al.<sup>125</sup> | https://github.com/VIB-PSB/i-ADHoRe |
| wgd v2 | Chen et al.<sup>126</sup> | https://github.com/heche-psb/wgd |
| divDyn v0.8.3 | Kocsis et al.<sup>114</sup> | https://cran.r-project.org/web/packages/divDyn/ |
| SciPy v1.10.1 | Virtanen et al.<sup>127</sup> | https://scipy.org/ |
| Python v3.8.3 | Python Software Foundation | https://www.python.org/ |
| MATLAB R2024a | MathWorks | https://www.mathworks.com/ |

### METHOD DETAILS

#### Collection and quality assessment of genome assemblies

We searched for all published angiosperm genome assemblies on the platform PlabiPD (January 5^th^, 2024). We downloaded at least one representative genome for most genera via the link provided in the data availability section of the corresponding article, leading to 470 genomes in total. To assess the quality of the genome assemblies and annotations, we used BUSCO (v5.4.3)^35^ (‘protein’ mode, embryophyta_odb10 database) and OMArk^36^ (Viridiplantae.h5 database).

#### Phylogenetic inference of angiosperms

Reconstructing species trees from gene trees involves uncertainties and biases such as incomplete lineage sorting (ILS) and sampling, horizontal gene transfer (HGT), gene flow, etc. Inferring orthologues de novo from a sequence similarity graph, as implemented in some ortholog inference algorithms^128,129^, is sensitive to parameters such as the inflation factor^130^, especially for single-copy orthogroups (SOGs). To avoid such biases, we used the “reference orthologues” from the embryophyta_odb10 BUSCO dataset^37^ to retrieve the orthologues in each one of the 470 angiosperm genomes, which were used to build individual gene trees, and summarize the species tree under the multispecies coalescence (MSC) model. Briefly, for each SOG in the *embryophyta_odb10* database, we first used HMMER (v3.1b2)^119^ to find corresponding orthologues using the pre-built profile hidden Markov model and the cut-off table from BUSCO. When multiple hits occurred, only the best hit was used for subsequent analysis. Secondly, a multiple sequence alignment (MSA) of nucleotides was inferred for each SOG using MAFFT (v7.310)^120^. IQTREE (v1.6.5)^121^ was then applied to infer the maximum likelihood (ML) gene tree for each MSA with the automatic detection of the best-fit substitution model via ModelFinder^122^. The uncertainties of branches were measured by the SH-aLRT^131^ and aBayes test with 1000 ultrafast bootstrap replicates, which were further optimized by a hill-climbing nearest neighbor interchange (NNI) search.

To find the optimal branch support cut-off to use for filtering branches, we tested cut-offs of 0%, 10%, 20%, 30%, 40%, 50%, 60% and 70%. To account for the effect of different MSA data types on the branch length and topology of inferred gene tree^132^, we conducted the gene tree inference using i) codon-level nucleotide; ii) 1^st^- and 2^nd^-position codon nucleotide; iii) TAPER-masked^53^ nucleotide; and iv) raw nucleotide datasets. Thus, we generated 32 gene tree datasets (eight branch support cut-offs x four MSA data types), each containing gene trees inferred from 1614 SOGs. The 1614 individual gene trees of each dataset were then supplied to ASTRAL-Pro^123^ to summarize the species tree under default parameters (Supplementary Information 1). We adopted the phylogenetic inference from the raw nucleotide dataset with the branch support cut-off of 50% based on its concordance with the statistical topological test and the optimized trade-off between high branch support of gene trees and high statistical support of the inferred phylogeny in terms of local posterior probability (LPP) (Supplementary Note 3).

#### Divergence time tree inference of angiosperms

Using the species tree obtained with ASTRAL-Pro, we conducted molecular dating analyses based on the concatenated amino acid MSA of 12 SOGs that covered at least 466 species (species coverage 99%) to obtain a time-calibrated tree. The time tree was inferred using the uncorrelated lognormal (UCLN) relaxed molecular clock model implemented in mcmctree (v4.9j)^124^, with soft fossil calibration bounds. The model parameters in mcmctree were set as follows: the clock model was set to ‘independent rates model’; a log-normal distribution of evolutionary rates across branches was set with a LG amino acid substitution matrix and a gamma model with five rate categories and α = 0.5; parameters for the birth-death process were set as 1 1 0.1 to generate uniform age priors on nodes that did not have a fossil calibration; gamma priors for the transition/transversion rate ratio and the shape parameter for variable rates among sites were set as 6 2 and 1 1; a Dirichlet-gamma prior was set with the parameters controlling the mean rate across loci and the logarithmic variance as 2 20 1 and 1 10 1. The first 2000 generations were discarded as burn-in after which 2,000,000 generations were pulled with sampling per 100 generations. To ensure accuracy and consistency, we assembled an angiosperm fossil calibration dataset (Supplementary Note 4) and the full age constraints implemented in our molecular clock analysis are shown in the calibration map (Supplementary Information 2). The effective sample size (ESS) was higher than 200 for all parameters, suggesting good convergence. Given the inferred time tree, the nucleotide substitution rate (Figure S2 in Document S1) was further estimated based on the concatenated nucleotide MSA of the same 12 SOGs using the best-fit substitution model predicted by ModelFinder^122^.

#### Identification and dating of putative WGD events

Intragenomic collinear segments and a burst of genes that seemed duplicated at the same time can be indicative of a putative WGD event. Here, we used i-ADHoRe (v3.0.01)^125^ and wgd v2^126^ to search for genomic collinearity and for bursts of gene duplication events in *K*_S_-based age distributions, where the number of duplicated genes is plotted against their age^105^. In short, the program “wgd dmd” was applied with default parameters to construct the whole paranome (collection of all duplicated genes in a genome) and the program “wgd syn” was applied with default parameters to infer genomic collinearity using i-ADHoRe. The whole paranome and ‘anchor pair’ (pairs of duplicated genes derived from large-scale gene duplication events and residing in duplicated segments) *K*_S_ distributions were then constructed using the program “wgd ksd” with default parameters^126^. We applied exponential-lognormal mixture models (ELMM) and log-scale Gaussian mixture models (GMM), as implemented in the program “wgd viz”, to delineate potential WGD components from the whole paranome and anchor pair *K*_S_ distributions, respectively. The phylogenetic location of putative WGD events can be inferred by ranking the relative timing of WGD events and speciation events within a *K*_S_ timeframe after proper correction of substitution rate variation^133^. The former are represented as peaks of the delineated log-normal components in the whole paranome and anchor pair *K*_S_ distributions while the latter are represented as peaks of the kernel density estimate (KDE) of the *K*_S_ distributions for orthologous gene pairs, identified as the reciprocal best hits (RBH) across all species.

For each putative WGD event, we compared its relative timing to a sequence of speciation events based on prior investigations of earlier studies to delineate its phylogenetic location (detailed species composition for each putative WGD event in Supplementary Note 6). The absolute dating of the identified WGD event was conducted in a standardized way. In short, given an identified WGD event for a group of species belonging to a certain order, we assembled an orthologous gene family dataset consisting of the anchor pairs and their orthologues from species of other orders (detailed order composition and species composition for each WGD event in Supplementary Note 6), in total 16-18 species per orthologous gene family following Chen et al. (2024)^126^. Based on the constructed orthologous gene family and our main tree topology, we conducted phylogenetic WGD dating on the concatenated amino acid MSA using mcmctree (v4.9j) with the same composition of ordinal age constraints and the same parameters as implemented in the divergence time tree inference section, except that the number of generations was increased to 20,000,000 and the sampling frequency to 1,000. For each WGD event, we dated at least one species, and we adopted the consensus mean, calculated as the overall mean of all WGD age estimates for a WGD event, and the 90% Highest Convergence Region (HCR), calculated as the intersection of the 90% Highest Posterior Density (HPD) of the WGD date estimates across species for a WGD event, to represent the age and the associated uncertainty of a WGD event. The estimated *K*_S_ peak and WGD date for each independent WGD event are summarized in Table S6.

#### Diversity dependence of WGD establishment rate and the deviation

The WGD establishment rate, measured by the number of established ancient WGDs per lineage per million years (My), likely covaries with the number of lineages, as ecological factors such as competition and niche overlapping are known to impact the establishment of polyploidy^10^. We applied a moving-window method to estimate WGD establishment rates over time. We found a significant linearity between the estimated WGD establishment rate profile and the number of lineages in the 466-species tree over time (*R*^2^ varying from 0.2734 to 0.8217 with varying window size, P<0.0001; Figure S3 in Document S1) and we showed that a linear model fit better than a power-law model in terms of AIC and BIC (window sizes starting from 8 My to prevent singularities derived from log-transformation; Figure S4 in Document S1), supporting a linear dependence of the WGD establishment rate on species richness. As the number of lineages grows exponentially in time (*R*^2^=0.9915, P<0.0001; Figure S5 in Document S1), the relationship between the WGD establishment rate and time is approximately an exponential of the form a.exp(bt)+c with a<0 and b,c>0. Substantial positive deviations of the WGD establishment rate profile from this relationship may indicate peaks in WGD establishment activity related to other (geological, environmental, biological) phenomena. To identify such peaks, a linear model was made of the WGD establishment rate moving average (window size 4 My) as a function of time and the number of lineages through time. The resulting relative residuals (observed data minus linear model estimates, normalized on the linear model estimates) are depicted in Figure 3A. Peaks in the relative residuals profile were identified using the ‘findpeaks’ function in MATLAB R2024a on a smoothened relative residuals profile (LOESS fit using the ‘smooth’ function in MATLAB R2024a with span set to 10% of the total number of data points).

#### Leave-one-out cross-validation for optimal KDE bandwidth

As an alternative to the moving-average analysis, we used kernel density estimation (KDE) to estimate the WGD establishment rate profile. To ensure proper smoothness while avoiding overfitting, we used leave-one-out cross-validation to find the optimal KDE bandwidth to model the WGD establishment rate over time. We compared 200 bandwidths evenly distributed from 1 to 20 My based on the average log-likelihood per test point. WGD age data was reflected around 0 Ma and the root age of the phylogenetic tree before KDE to avoid boundary effects. We found that 1) starting from bandwidth 3.1, the increase in average log-likelihood between adjacent bandwidths significantly slowed, i.e., became smaller than 5% of the total range of average log-likelihoods for 10 consecutive steps; 2) after bandwidth 10.1, the average log-likelihood declined monotonically, as shown in Figure S8 in Document S1. Given the marginal improvement of log-likelihood in between, we used a lower bandwidth (3.1) that gave higher resolution to detect WGD peaks. The resulting KDE curve was converted to a curve of WGD establishment rate per lineage by dividing the kernel density estimates by the number of lineages in the species tree at each time point.

#### Exponential-Gaussian mixture model of WGD establishment rate

To assess how many peaks occur in the WGD establishment rate profile obtained through KDE, we fit exponential-Gaussian mixture models with an exponential background and 6 to 10 Gaussian components to the profile using the CurveFitter toolbox in MATLAB R2024a. An exponential background of the form a.exp(bt)-a with a<0 and b>0 was used to model the observed lineage diversity dependence of the WGD establishment rate (with the term -a used to force the background to go through the origin at 0 Ma). The highest adjusted *R*^2^ value was achieved for the model with 9 Gaussian components, including one underlying a non-visible peak at ∼66 Ma that is masked by the peaks before and after it.

#### Homogeneous Poisson process model of WGD simulation

Inspired by Vanneste and co-authors^24^, we devised a basic null model of WGD simulation in Python 3.8.3 whereby WGD events along each branch of the time-calibrated phylogenetic tree were modeled as an independent homogeneous Poisson process with the time interval between events exponentially distributed and controlled by a WGD establishment rate parameter. To accommodate potential variation of WGD establishment rates across different clades, we allowed clade-specific rate parameters for the ANA grade, Ceratophyllales, Chloranthales, eudicots, monocots and magnoliids, respectively. We calculated the clade-specific rate parameters that maximized the likelihood given the independent homogeneous Poisson process model and the observed WGD events as the phylogenetic background rate parameters for each clade (Table S7). We also validated that this variable-rate model fit the data significantly better than the alternative simpler model with a constant rate across all clades (likelihood ratio test, P <0.0001). Under this independent homogeneous Poisson process model and the clade-specific background rate parameters, we simulated WGDs throughout the main tree and documented the randomly generated WGD ages for 10,000 replicates to construct the null distribution. KDE profiles of the WGD establishment rate were constructed for each set of simulated WGD ages as described above, using the same bandwidth (3.1). The mean, median, 50%CI, and 95%CI of the simulated WGD establishment rate profiles were calculated over time and compared to the observed WGD establishment rate profile.

#### Non-homogeneous Poisson process model of WGD simulation

As the WGD establishment rate is related exponentially to time because of the observed negative linear lineage diversity-dependence of WGD establishment and the exponential increase of lineages over time, we devised a non-homogeneous Poisson process (NHPP) model to build an alternative null distribution of WGDs accounting for the WGD establishment rate variation across time using a standard thinning method. In this model, WGDs are generated following a Poisson process for each phylogenetic branch with exponentially decaying rate from the root to the tips of the tree. The WGD establishment rate is assumed to follow the functional form a.exp(bt) + c over time t, with parameters a < 0, b, c > 0. We first applied a global optimization approach using the differential evolution algorithm (maximum 10,000 iterations) from the SciPy library in Python 3.8.3 to estimate the parameters a,b and c of the NHPP model based on the observed WGD profile. We varied the random seed as an integer from 0 to 9,999 and repeated the optimization process 10,000 times, the median estimate of which was then integrated into the WGD establishment rate function for simulation, i.e., a = -0.01506, b = 0.01949, c = 0.01538. Subsequently, 10,000 NHPP model simulations were performed and KDE profiles of the WGD establishment rate were constructed for each set of simulated WGD ages as described above, using the same bandwidth (3.1). The mean, median, 50%CI, and 95%CI of the simulated WGD establishment rate profiles were calculated over time and compared to the observed WGD establishment rate profile. We also increased the number of simulations to one million and obtained nearly identical results, confirming the sufficiency of our simulation.

#### Fossil-based genus-level extinction rate estimation

We downloaded the latest angiosperm fossil occurrence data from the Paleobiology database^113^ on July 10th 2025, for the time frame spanning the Triassic until now. We circumvented potential homonymies by combining the class names and genus names to create individual entries. Multiple species of the same genus that were registered in the same collection are only counted as one occurrence. To take into account variations derived from different methodologies, we implemented three methods of sampling standardization (i.e., raw pattern, Classical Rarefaction^134^ and Shareholder Quorum Subsampling^135^) and four macroevolutionary rate models (i.e., per capita rates^136^, corrected three-timer rates^137^, gap-filler equations^138^ and second-for-third substitution rates^139^) using divDyn (v0.8.3)^114^ to estimate angiosperm genus−level extinction rates based on the time-scale of geological stages^140^.

#### Statistical tests for association between WGD establishment rate peaks, geological events, extinction events and paleoclimatic extrema

Nine peaks in the relative residuals of the linear model for the WGD establishment rate as a function of time and the number of lineages were supported by Gaussian components in the optimal exponential-Gaussian mixture model. Associations between these peaks and geological events, genus-level angiosperm extinction events and extrema in paleoclimatic variables, as depicted in Figure 3, were assessed through two types of Monte Carlo randomization analysis. In the first set of analyses, for each data layer the WGD peaks are compared to, 9 peak ages were randomly sampled a million times from a uniform distribution on the interval [0 Ma, 115 Ma]. For each random sample, the average nearest distance of the sampled WGD peak ages to the ages of ‘events’ in the other data layer were computed. Event ages are defined as the mean ages for geological events, the ages of maxima in the genus-level angiosperm extinction rate curve, and the ages of maxima and minima in the other data layers. Average nearest distances were computed with respect to the peaks in the other data layer, not the WGD peaks. For the paleoclimatic profiles exhibiting many peaks of varying peak importance (δ^13^C, δ^18^O, CO_2_ sea surface temperature, global mean surface temperature), only the top-18 minima and maxima in terms of peak importance were taken into account, and association analyses were done for different peak sets containing the top-x peaks with x=1. 18. For each peak set tested, the average nearest distance of the observed WGD peak ages to the peak set was compared to the sampled average nearest distances under the null hypothesis of uniformly distributed peak locations, and a P-value for peak association was computed as P=(k+1)/(n+1) with k as the number of times the sampled average nearest distance is lower than or equal to the observed average nearest distance and n as the number of random samples (1 million)^141^.

The second set of randomization analyses and P-value calculations was performed in the same way, except that distances between WGD peaks were sampled instead of WGD peak ages. For each random sample, an age in the interval [0 Ma, 30 Ma] was sampled to position the youngest random WGD peak, and subsequent peaks were positioned at inter-peak distances sampled from a Gaussian distribution fit to the inter-peak distances for the 9 observed WGD peaks (μ=11.57, σ=3.37 Ma).

All association analyses were done in MATLAB R2024a. Maxima were computed using the ‘findpeaks’ function on the raw time profile (for the genus-level angiosperm extinction rate, global mean surface temperature and CO_2_ concentration), a smoothing spline (smoothing parameter 0.05 for δ^13^C and δ^18^O isotopic signature profiles) or a LOESS fit (with span = 2% for the sea surface temperature data). Minima were computed using the ‘findpeaks’ function on the opposite (additively inverted) profiles.

### QUANTIFICATION AND STATISTICAL ANALYSIS

In the statistical test for the support of competing topologies pertaining to Chloranthales, we conducted one-way ANOVA and Tukey’s HSD test to determine which topology gets significantly higher support among different datasets, details in Supplementary Note 3. In the statistical test for the support of competing topologies pertaining to Crossosomatales, we conducted two-sided t-test and Mann-Whitney U test to determine which topology gets significantly higher support among different datasets, details in Supplementary Note 3. In the delineation of random sampling bias, we conducted ordinary linear regression to demonstrate the significantly reduced random sampling bias of our main tree, details in Supplementary Note 3. To compare our divergence time estimates with other studies, we calculated the Pearson correlation coefficients of the estimated stem order and family ages between our main tree and that of Ramírez-Barahona et al. (2020)^43^, details in Supplementary Note 5. Relative nucleotide and amino acid substitution rates across angiosperms were compared by one-way ANOVA and Tukey’s HSD test, details in Supplementary Note 5. Significance is defined by a P-value lower than 0.05, as denoted alongside the associated statistical tests.

### ADDITIONAL RESOURCES

Description: https://bioinformatics.psb.ugent.be/AngioWGD/

To adhere to the FAIR data principles (Findable, Accessible, Interoperable, and Reproducible), we developed the *AngioWGD* website which can be used to visualize and download data on WGD events across the plant phylogeny.

