## Supplementary Materials for "The Rise of Polyploids During Environmental Upheaval"

### Supplementary Note 1 The *AngioWGD* web application

To ensure all data generated in this work abide by FAIR data principles (Findable, Accessible, Interoperable, and Reproducible), we developed *AngioWGD*, a user-friendly R Shiny application that can be used to visualize and download data on WGD events across the plant phylogeny. The application is available online at https://bioinformatics.psb.ugent.be/AngioWGD/, and it is distributed as an R package (available at https://github.com/almeidasilvaf/AngioWGD) that can be installed locally, ensuring users can always access the application, even in case of server downtime. The app comprises three main pages, namely 'Explore WGD events', 'WGDs by species', and 'Original data'.

The 'Explore WGD events' page can be used to search for WGDs in particular angiosperm clades (e.g., families, orders, and larger clades such as 'Monocots' and 'Fabids'), and visualize a phylogenetic tree with WGD events placed on nodes, as well as an interactive table with date statistics and phylogenetic location for each WGD event (Figure S1a). The 'WGDs by species' page can be used to explore an interactive table of WGD events for each species, including $K_S$ peaks, WGD ages, and uncertainties around point estimates, and to visualize posterior distributions of WGD ages for selected WGD events (Figure S1b). Finally, the 'Original data' page contains taxonomic information for all species used in this study, and links to seamlessly download coding sequence (CDS) and annotation files (.fasta and .gff3, respectively) stored in FigShare repositories associated with this work. All figures and tables in the app can be saved to files (.svg and .pdf for figures, .tsv for tables) for downstream analyses and/or to be included in publications.

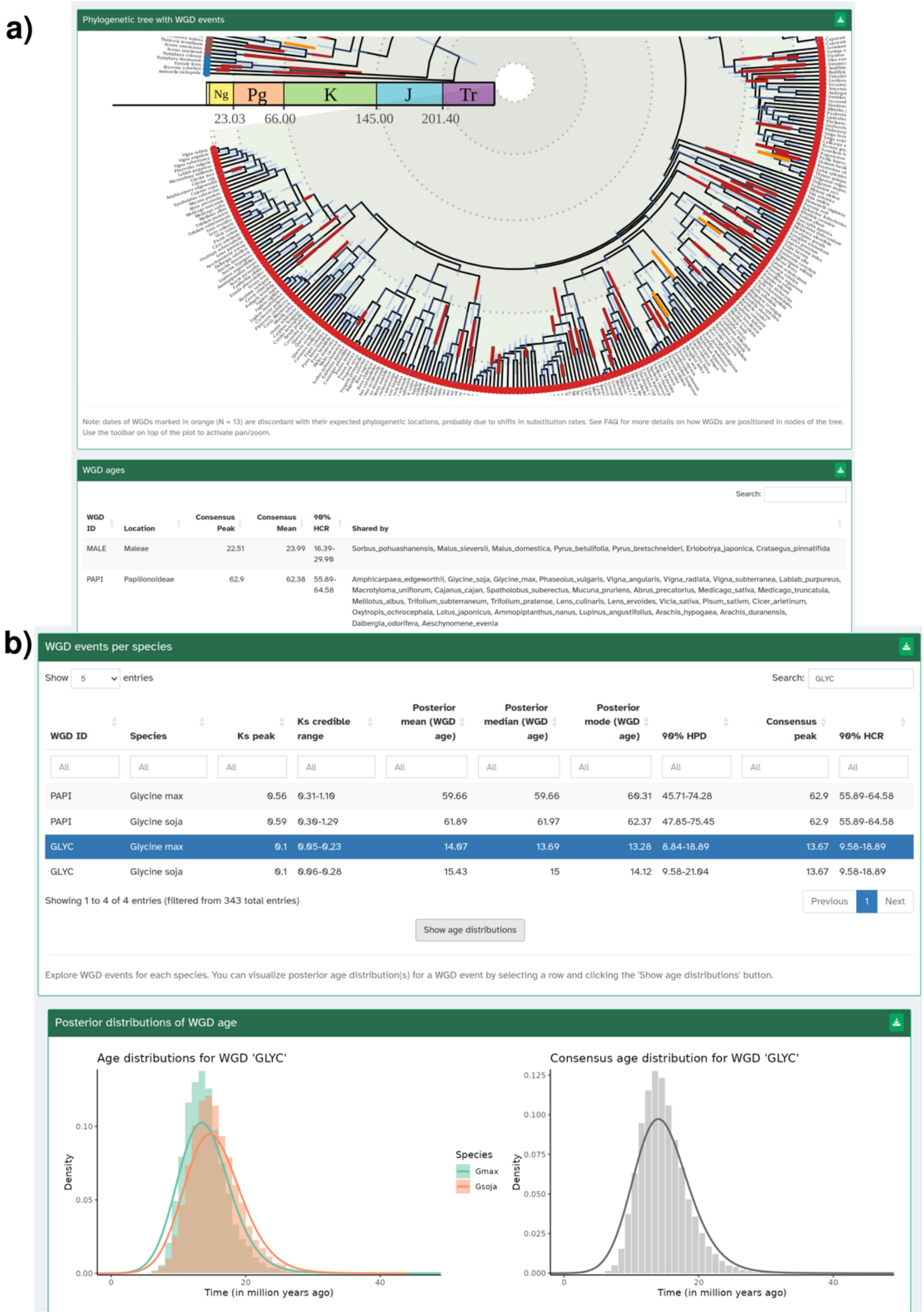


**Figure S1 | Main functionalities of the *AngioWGD* web app.** a) Use case for the 'Explore WGD events' page. The interactive figure (top) shows a zoomed-in phylogenetic tree. Rectangles in tree nodes indicate WGD events, with rectangle widths indicating the 90% HCR of WGD dates. The interactive table (bottom) shows the dates for all WGD events in the selected clade (or clades by default). b) Use case for the 'WGDs by species' page. The interactive table (top) shows WGD date statistics ($K_S$ peak, WGD dates, and uncertainties around estimates) for each species after searching for the text 'GLYC'. Upon selection of a particular WGD event ('GLYC' in this example, representing the WGD event in the common ancestor of *Glycine* sp.), users can visualize posterior distributions of WGD ages for each species separately (if the selected WGD was identified in multiple species), and for all species combined (i.e., the 'consensus' distribution).

**Supplementary Note 2 Assessment of genome assembly**

Although new genomes are being sequenced and assembled at an unprecedented rate, there is no systematic assessment of the quality of such bulk data, which is essential for the downstream comparative genomic analysis. Here we assessed our large genome dataset composed of 470 different angiosperm species and 1 outgroup species *Cycas panzhihuaensis* using BUSCO[1] (v5.4.3) and OMArk[2]. The database embryophyta_odb10[3] and the proteins mode were used for BUSCO. The database Viridiplantae.h5 was used for OMArk.

As presented in Figure S1, the mean, median, mode, Q1 and Q3 of the complete gene score evaluated by BUSCO were 87.71%, 92.00%, 94.59%, 85.15% and 96.45%, respectively, indicating that most of the genome assemblies were of good completeness. The mean, median, mode of the missing gene score were 7.93%, 4.10%, 3.21%, respectively, suggesting a generally low percentage of missing genes in most of the genome assemblies. The mean, median, mode of the fragmented gene score were 4.36%, 2.70%, 1.74% respectively, suggesting an even lower percentage of fragmented genes in most of the genome assemblies. The histogram of the single gene score showed a rather different pattern than the last three scores that multiple salient peaks emerged, suggesting distinct process of single-copy gene retention across different lineages. The histogram of the duplicated gene score also showed distinct gene duplication dynamics of different lineages that while the median and mode were low as 7.10% and 4.81%, the high mean as 18.50 implied that there were some genomes bearing remarkably high percentage of duplicated genes. As shown in Figure S2, the mean, median, mode, Q1 and Q3 of the complete gene score evaluated by OMArk were 92.15%, 94.45%, 95.62%, 90.41%, 96.52%, respectively, generally higher than that of BUSCO, suggesting an overall high level of completeness of the genome assemblies. The mean, median, mode of the missing gene score were 7.85%, 5.55%, 4.38%, respectively, suggesting a low level of missing genes in most of the genome assemblies. The histogram of the single and duplicated gene scores were analogous to that of BUSCO in which multiple conspicuous uplifts emerged, suggesting disparate pattern of duplicated and single-copy gene retention across lineages. The expected and unexpected duplicated gene were determined by whether the genes that were found in multiple copies corresponded to a known duplication event in descendants of the reference gene family or not[2]. The mean, median, mode of the expected duplicated gene score were 5.03%, 1.20%, 1.26%, respectively, suggesting that most of the duplication events were indeed also found in the precomputed reference gene families of the Viridiplantae.h5 database for most of the genome assemblies. The consistency assessment of OMArk provides the proportion of the genes in the query genome that is likely corresponding to an actual gene by comparing to the known gene families of the selected ancestral lineage[2]. The inconsistent gene score measures the percentage of genes that are corresponding to known gene families but from different lineages in a seemingly random manner. The unknown gene score measures the percentage of genes that do not share enough similarity with known gene families and are as such likely orphan genes. OMArk infers a main species or clade that is most consistent with the taxonomic distribution of the gene families where the genes from the query genome were assigned and the percentage of associated queried genes is provided as the so-called “main species-consistent score”. The mean, median, mode of the consistent gene score were 82.83%, 84.97%, 90.11%, respectively, suggesting that most of the genes were presented in the ancestral gene repertoire already for most of the genome assemblies. The mean, median, mode of the unknown gene score were 14.75%, 11.72%, 7.42%, respectively and we also observed a multimodal distribution, suggesting distinct dynamics of novel gene acquisition across lineages. The mean, median, mode of the main species-consistent score were also high as 85.25%, 88.28%, 92.58% respectively, suggesting high consistency with existing genomes. Altogether, the assessment of BUSCO and OMArk showed that the genome assemblies of our dataset have generally high completeness and consistency.

Furthermore, we conducted a comparison between the evaluation from BUSCO and OMArk. As shown in Figure S3, there were significant correlations between the results of BUSCO and OMArk (p-values all smaller than 0.0001 of both Pearson correlation test and linear regression analysis), in particular on the single gene score and duplicated gene score with Pearson correlation coefficients as 0.8074 and 0.8199, suggesting that the assessment of the gene copy state might be more conservative than that of completeness or missing (Pearson correlation coefficients as 0.5382 and 0.6144), which is largely dependent upon the amount of adopted genomes.

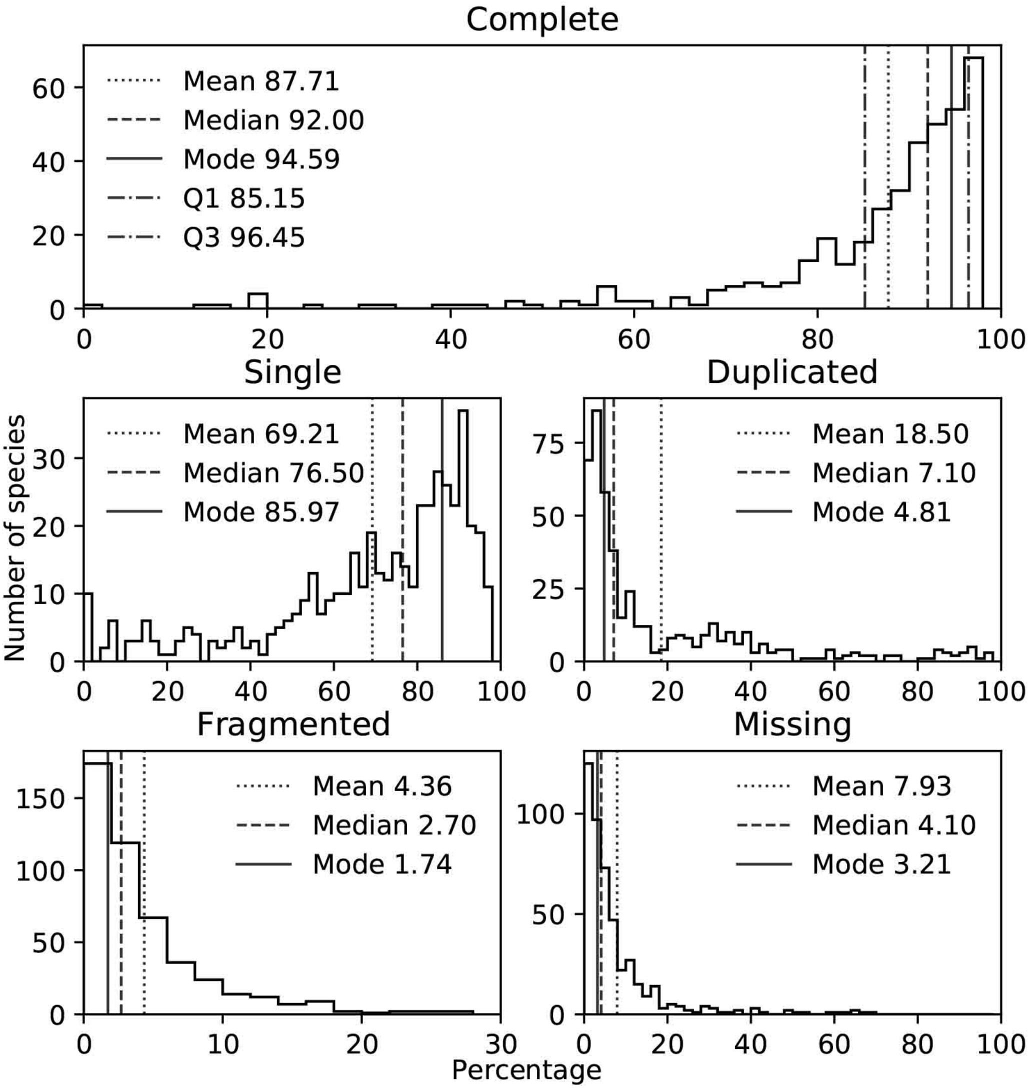


**Figure S1 | Genome assessment based on BUSCO.** The scores assessed by BUSCO for the complete, single, duplicated, fragmented and missing genes are presented in different panels. The mean, median, mode, Q1 and Q3 of the complete gene score and the mean, median and mode of the other scores are denoted.

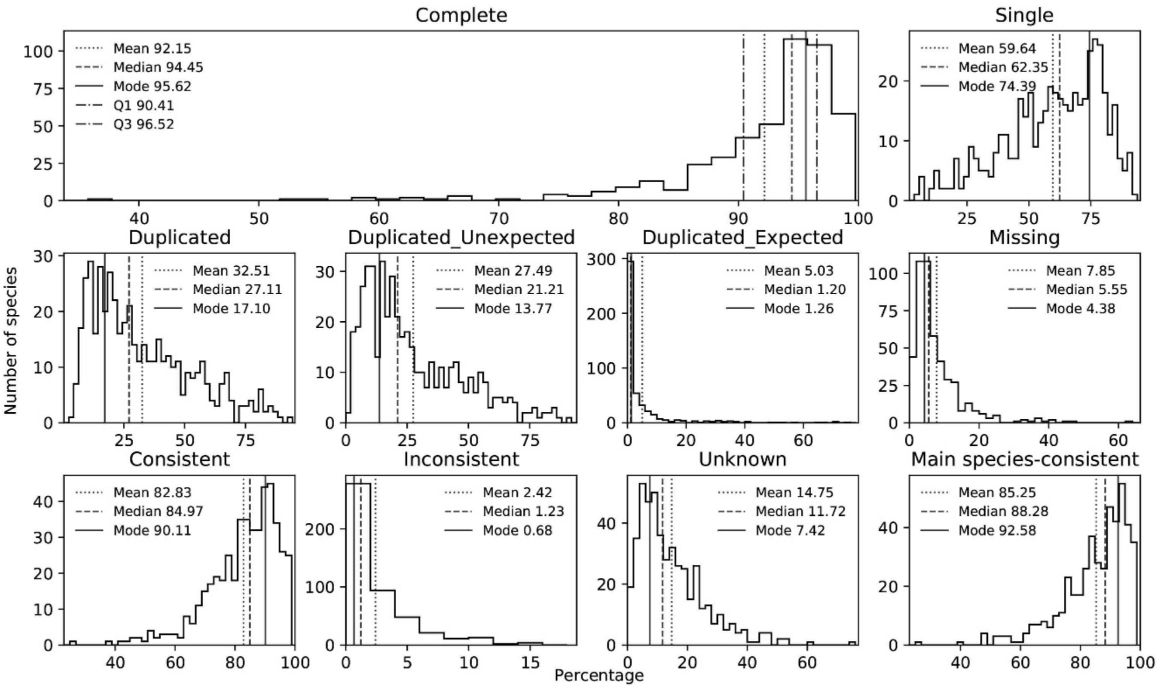


**Figure S2 | Genome assessment based on OMArk.** The scores assessed by OMArk for the complete, single, duplicated, duplicated and unexpected, duplicated and expected, missing, consistent, inconsistent, unknown, and main species-consistent genes are presented in different panels. The mean, median, mode, Q1 and Q3 of the complete gene score and the mean, median and mode of the other scores are denoted.

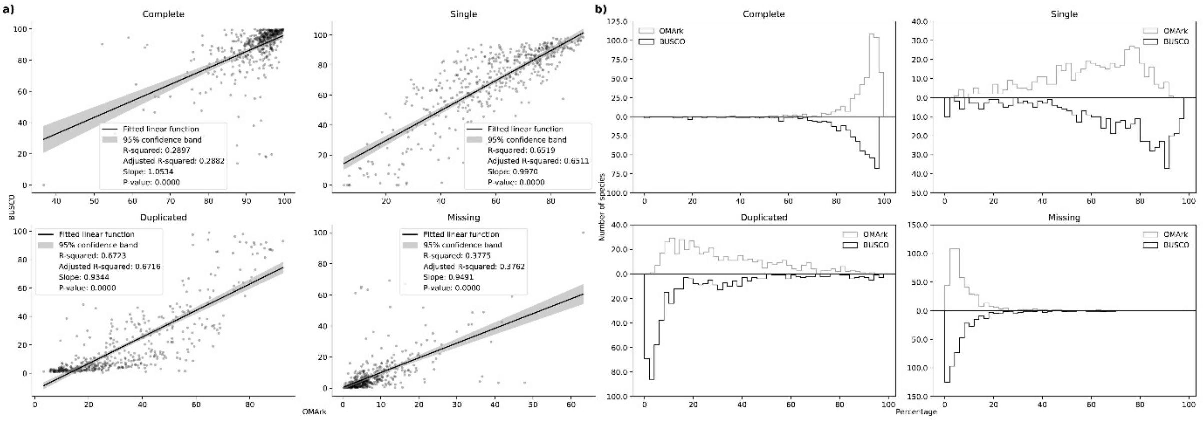


**Figure S3 | Comparsion between BUSCO and OMArk.** Panel a shows the linear correlation between different scores evaluated by BUSCO and OMArk with the 95% confidence band, R-squared, adjusted R-squared, slope, and p-value of the linear regression function denoted. Panel b shows the bihistogram of different scores evaluated by BUSCO and OMArk.
